# Quorum sensing and DNA methylation play active roles in clinical *Burkholderia* phase variation

**DOI:** 10.1101/2024.09.30.615881

**Authors:** Pauline M.L. Coulon, Marie-Christine Groleau, Abderrahman Hachani, Matthew P. Padula, Timothy P. Stinear, Eric Déziel

## Abstract

Phenotypic diversity in bacteria often results from adaptation to changing environmental conditions and is exemplified by variable colony morphotypes. Discrete genomic changes and modulation in gene expression occur in *Burkholderia pseudomallei* undergoing adaptation. Alternatively, adapted colony morphotype variants of species belonging to the *Burkholderia cepacia* complex (Bcc) lose a complete replicon (the pC3 virulence megaplasmid), which affects their production of virulence factors. We report that variants arising in *Burkholderia ambifaria* clinical isolates - with affected phenotypes - have retained their pC3, suggesting that another phase variation mechanism can take place in this Bcc species. Proteomic and phenotypic characterisation showed that morphotype variants of *B. ambifaria* strains CEP0996 (pC3-null) and HSJ1 (pC3-positive) share similarities in phenotypes controlled by the Cep quorum sensing system. Thus, we determined the role of quorum sensing in *B. ambifaria* HSJ1 phase variation and confirmed that the main quorum sensing system Cep is important for the emergence of variants. As DNA methylation is one of the main epigenetic factors involved in bacterial phase variation that regulates some virulence factors of the Bcc species *Burkholderia cenocepacia*, we hypothesized that *B. ambifaria* HSJ1 phase variation could also be regulated by adenosine DNA methylation. By deleting the three putative adenosine DNA methyltransferases, we found that an orphan type II DNA methyltransferase prevents the emergence of phase variants. This is the first study to report quorum sensing and adenosine DNA methylation as two antagonistic systems independently controlling phase variation.

**Importance:** Some *Burkholderia* species are pathogenic to plants, animals, or humans. In immunocompromised individuals, such as those with cystic fibrosis, infection with *Burkholderia cepacia* complex (Bcc) bacteria can lead to "*cepacia* syndrome." In the Australian Aboriginal population, melioidosis caused by *B. pseudomallei* is prevalent, particularly among those with diabetes or alcoholism. *Burkholderia*’s phenotypic plasticity, including colony morphotype variation (CMV), enables rapid adaptation to diverse environments, enhancing survival and pathogenicity. This study reveals phase variation as a new CMV mechanism within the Bcc group. We found that quorum sensing and DNA methylation are involved in phase variation. Understanding the underlying CMV mechanisms could lead to the development of targeted therapies against these highly antibiotic-resistant bacteria.

## Introduction

The *Burkholderia* genus encompasses both plant-beneficial and pathogenic species (1). Some species such as *Burkholderia glumae* and *Burkholderia gladioli* are well described plant pathogens, responsible for bacterial panicle blight or leaf strike (2–5). The “*Bptm*” group includes the pathogens *Burkholderia pseudomallei* (*Bp*) and *Burkholderia mallei*, causing melioidosis in humans and glanders in equines, respectively (6–8). The *Burkholderia cepacia* complex (Bcc) comprises species that are broadly distributed in a variety of environments such as soils, water, or plant rhizosphere (9). This group has emerged as opportunistic pathogens that can cause the ‘*cepacia* syndrome’ in people with cystic fibrosis (CF) and immunocompromised individuals (1, 10–12). However, some Bcc members also have bioremediation and plant growth-promoting properties (e.g. *B. ambifaria* (*Ba*) and *B. vietnamiensis*) (13–16) or are plant pathogens (e.g. *B. cepacia*) (17); reviewed by L. Vial et al. (18). Understanding *Burkholderia* virulence and persistence during infection is essential due to their high intrinsic tolerance and resistance to clinically relevant antibiotics (19–21).

Phase variation is mostly a reversible process of environmental adaptation (22, 23) due to (i) genomic variations (e.g.: SNP, indels or sequence inversion) (24), (ii) modulation of gene expression (25–29) or (iii) epigenetic factors such as DNA methylation (22, 30, 31). It produces phenotypic changes that can promotes bacterial fitness. Phase variation has been described in *Burkholderia* species and observed in pure cultures and in samples from long-term CF infections (23, 24, 28, 32–39). In members of the Bcc, phase variation has been described among clinical *Ba* isolates; such variant colonies are readily recognized by their smooth and translucent phenotype, whereas wildtype colonies of clinical isolates are rough and wrinkled (23). We have previously reported clinical *Ba* strains that switched to a variant phenotype with characteristics mostly associated with environmental strains better adapted to the rhizosphere by displaying a reduction in virulence factors such alginate-like exopolysaccharide, hemolysin production, antifungal and cholesterol oxidase activities and 4-hydroxy-3-methyl-2-alkylquinolines (HMAQ) production (23). Bernier et al. (26) have shown that shiny variant colonies appear from the clinical *Bc* strain K56-2 due to a mutation in the LysR-type transcriptional regulator ShvR. Other than colony morphotype, antimicrobial, protease and exopolysaccharide (EPS) production as well as biofilm formation are also controlled by ShvR (26, 27, 40–42). Using a Tn*5* random mutagenesis library in the clinical *Bc* strain H111, Agnoli et al. (32) discovered that multiple non-pathogenic shiny colony clones were lacking their third replicon, described as a megaplasmid of virulence (pC3). Following curing of the pC3 from six Bcc strains, virulence (in *Galleria mellonella* and *Danio rerio* infection models) was attenuated and the PC3-null derivatives showed diminished antifungal activity, EPS and protease productions, and altered substrate utilization (32, 43, 44). Although the loss of pC3 has been described as an adaptation to laboratory conditions, four environmental *B. ubonensis* isolates were found as having irreversibly lost their pC3 (45), indicating that this phenomenon can occur naturally.

Recently in the *Bc* strains J2315 and H111, the pC3 was shown to be stabilized by DNA methylation, as deletion of adenosine DNA methyltransferases increased the occurrence of variants lacking the replicon (46). DNA methylation is used as an ON/OFF epigenetic regulation system by methylating or demethylating specific motifs in the promoter region of transcriptional regulators and genes coding for virulence functions such as biofilm formation, cell aggregation and motility in *Bc* (47). Such virulence factors are also controlled by quorum sensing (QS), an intercellular communication system dependant on cell-cell signalling to synchronize target gene transcription within bacterial populations (48). In members of the Bcc, QS is typically mediated by LuxI/LuxR type systems (49). Each system is comprised of one acyl-homoserine lactone (AHL) synthase (CepI) and one cognate transcriptional regulator (CepR). The AHLs produced by CepI bind specifically to CepR, forming an activated CepR/AHL autoregulatory complex controlling the expression of *cepI*, *cepR* and several target genes, including ones encoding for virulence factors (50–54). In *Bc*, AHL QS interplay with other QS regulatory elements such as the *Burkholderia* diffusible signal factor (BDSF) system which inhibits the production of cyclic-di-GMP and thus the biofilm formation and the transcription of *cepI* (55–59), and activation of ShvR, a transcriptional regulator affecting the expression of over a thousand genes, including 263 genes co-regulated by QS such as *cepI*, *cepR* and *cciI/cciR* (a second AHL-based QS system) (26, 27, 40–42).

In this study, we report that our previously described phase variants of clinical *Ba* strains (23) actually belong to two groups: (i) pC3-positive phase variants; and (ii) pC3-null phase variants.

## Results

### Clinical *B. ambifaria* strains generate two distinct phenotypic variant types, including one which loses its pC3 replicon

HSJ1 is a clinical *Ba* strain that generates a phenotypic variant distinguishable on agar plates containing Congo Red (23). One characteristic of the variant is the loss of production of secondary metabolites belonging to the 4-hydroxy-3-methyl-2-alkylquinoline (HMAQ) family produced by enzymes encoded by the *hmqABCDEFG* operon, which is normally located on the pC3 replicon (23, 60, 61). We previously established the prevalence of the Hmq system in the Bcc (61) using PCR primers specific to the first and the last gene of the *hmq* operon (namely *hmqA* and *hmqG*). Using the same method, we found that the *hmq* genes were absent from two of the eight investigated clinical (**Table 1**). Interestingly, out of the six remaining strains carrying the *hmq* operon, the *hmqA* and *hmqG* genes were not detectable in four of their respective variants (**Table 1**). Together these results hinted to a possible loss of the pC3 operon in up to six out of eight *Ba* clinical strains.

**Table 1.** Three-quarters of phenotypic variants derived from eight clinical *Ba* isolates have lost their pC3 replicon.

| Strains |  | *hmqA* | *hmqG* | *cepl2* | BAMB_RS28665 | BAMB_RS29710 | BAMB_RS31530 |
| --- | --- | --- | --- | --- | --- | --- | --- |
| *B. ambifaria*<br>CEP0516 | WT | - | - | + | + | + | + |
|  | Variant | - | - | ? | ? | ? | ? |
| *B. ambifaria*<br>CEP0617 | WT | + | + | + | + | + | + |
|  | Variant | ? | ? | ? | ? | ? | ? |
| *B. ambifaria*<br>CEP0958 | WT | + | + | + | + | + | + |
|  | Variant | ? | ? | ? | ? | ? | ? |
| *B. ambifaria*<br>CEP0996 | WT | + | + | + | + | + | + |
|  | Variant | ? | ? | ? | ? | ? | ? |
| *B. ambifaria*<br>AU4157 | WT | + | + | + | + | + | + |
|  | Variant | ? | ? | ? | ? | ? | ? |
| *B. ambifaria*<br>AU8235 | WT | - | - | + | + | + | + |
|  | Variant | - | - | ? | ? | ? | ? |
| *B. ambifaria*<br>HSJ1 | WT | + | + | + | + | + | + |
|  | Variant | ✓ | ✓ | ✓ | ✓ | ✓ | ✓ |
| *B. ambifaria*<br>AU0212 | WT | + | + | + | + | + | + |
|  | Variant | ✓ | ✓ | ✓ | ✓ | ✓ | ✓ |

As some Bcc isolates have previously been reported to have lost their pC3 (32, 43–45), we performed a PCR screen for five additional genes located on the pC3 replicon (*cepI2*, BAMB_RS28665, and BAMB_RS31530). The gene *hisA*, located on chromosome 1 (C1), was used as a positive control and could amplify in both the parental and variant strains. As suggested by the initial *hmq* gene screen (61), no amplification was observed in six out of eight tested variants, whereas amplification was positive for their respective parental strains (**Table 1**). These results indicated that several *Ba* variants indeed lost the pC3 replicon and were identified as “pC3-null”. On the other hand, variants of strains HSJ1 and AU0212 were positive for all screened genes, including *hmqA* and *hmqG*, even though they do not produce HMAQs (**Table 1**; (23, 61)). The PCR results were confirmed by whole genome sequencing of the WT and variants of strains CEP0996 and HSJ1, representing both types of variants, using long and short-read sequencing (**Figure S1**). Indeed, by mapping each variant’s Illumina reads onto their respective parental strain’s whole-genome assembly, we confirmed that CEP0996v had indeed lost its pC3 (**Figure S1A**) while it was still present in HSJ1v (**Figure S1B**). CEP0996 and its pC3-null variant (CEP0996 pC3-null) as well as HSJ1 and its pC3-positive variant (HSJ1v) were chosen as models to further characterise phase variation in *Ba*.

### Both types of variants (pC3-null and pC3-positive) share similarities and dissimilarities in the production of QS-regulated factors

Since several phenotypes are in common between both types of variants, such as loss of HMAQ production (23), we sought to further characterize the different features of pC3-null variants compared to pC3-positive ones.

The absence of production of HMAQs suggested that some genes on the pC3 of HSJ1v are not expressed. We therefore compared the proteomes of both variants with their parent strains when cultured in TSB to an OD_600_ of ∼1 (**Figure 1**). For the CEP0996 strain, 2872 proteins out of 6508 open reading frames (ORF) encoding those proteins were quantified while 2850 proteins out of 6700 ORF encoding those proteins were quantified for HSJ1 (**Figure S2**). Based on our analysis, 396 proteins were present or absent in CEP0996 pC3-null compared to its WT. From those, 104 are encoded on chromosome 1, 112 on chromosome 2 and 180 on pC3 (**Figure 1A; Tables S6-7**). For HSJ1, 328 proteins were present or absent in the variant vs the WT, plus 20 proteins whose abundance was significantly different (with an FDR adjusted p-values below 0.01) between HSJ1 and HSJ1v. From those 348 proteins, 122 are encoded on chromosome 1, 125 on chromosome 2 and 101 on pC3 (**Figures 1A**; **Tables S8-9**). Among the 6461 homolog proteins between CEP0996 and HSJ1, 77 were significantly different in abundance in variants compared to their WT including 16, 28, 32 respectively encoded on chromosome 1, 2 and pC3; and one encoded on CEP0996 chromosome 1 but located on chromosome 2 in HSJ1.

**Figure 1:**
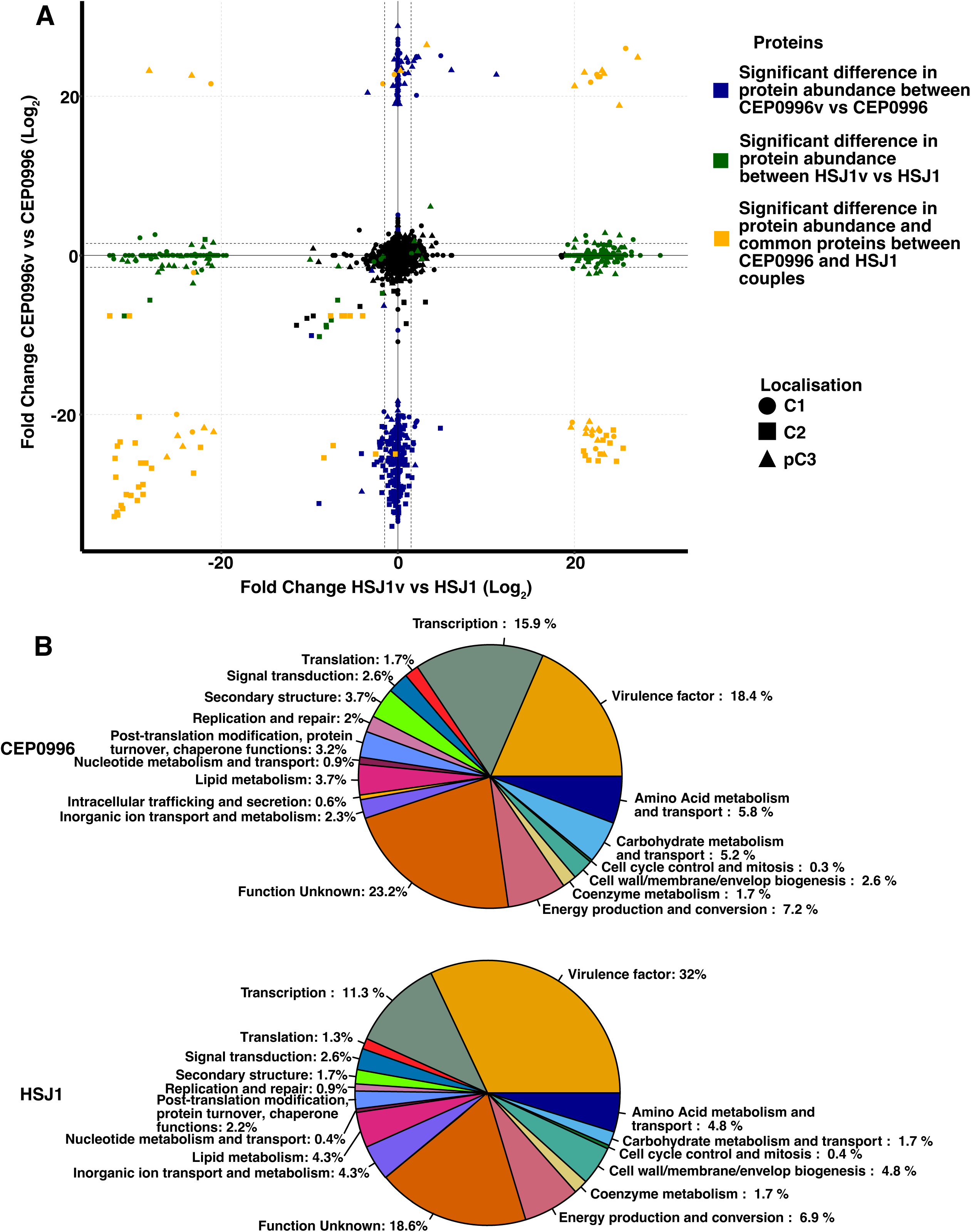
Both types of variant showed dissimilarities in protein production signatures. A) Proteome of both WT and variants for both CEP0996 and HSJ1. Significative differences in protein abundance between CEP0996v and CEP0996 are shown in blue, significative differences in protein abundance between HSJ1v and HSJ1 are shown in green, significant common proteins between CEP0996 and HSJ1 couples are shown in yellow. Proteins located on chromosome 1 are represented by circles, proteins located on chromosome 2 are represented by squares, proteins located on pC3 are represented by triangles. B) Significant proteins have been classified by “Clustering of orthologous groups” using Eggnote tool v5 (126). Proteins found in CEP0996 couple are in blue while Proteins found in HSJ1 couple are in green.

Overall, these proteins belong to primary metabolism and energy production, antibiotic biosynthesis clusters, and virulence factors (**Figure 1B; Tables S10-11**) – which was also reported to be affected by colony morphotype variation in *Bp* (62–65). Interestingly, type II-IV pilus biosynthesis, cepacian EPS, secondary metabolites biosynthesis enzymes, such as those responsible for the production of the AFC lipopeptide, enacyloxin, HMAQs, occidiofungin and pyrrolnitrin – all known to have antimicrobial properties (66–72), were reduced in abundance in both types of variants (**Table S11**). These data could explain the loss of antifungal activity previously reported for both type of variants (23). This could also suggest their reduced ability to compete in the CF lung environment, as well as altered Ba signaling and immune evasion due to a decrease in Hmq proteins—similar to what has been proposed in Bp for long-term infection (73). However, proteins linked to virulence factors – such as adhesins, proteins involved in flagellum biosynthesis, including the non-specific flagella regulator FlhD (74) and siderophores – were increased in abundance in the variants, corroborating the overproduction of siderophore in HSJ1v that we previously reported (23).

Following our proteomic analysis, we compared phenotypes between parental strains and their corresponding variants (23, 32). Colony morphologies on CRLA plates, biofilm production, swimming motility, flagella observation using electronic microscopy, and siderophore production were assessed (**Figure 2; Tables S12**). In bacteria, Congo red binds to different EPS such as cellulose or amyloid fibers (75, 76). While no homolog genes encoding for amyloid adhesin were found in both CEP0996 and HSJ1 isolates, our proteomic analysis revealed that proteins encoded by the bacterial cellulose synthase (*bcs*) cluster genes are similarly produced by WT and variants (77) (**Table S6 and Table S8**). Thus the difference in Congo Red binding by variant colonies (**Figure 2A**), could be explained either by the lack of production of the EPS cepacian production on CRLA (78) (**Table S11**), by a difference in other EPS production (79) such as galactan-3-deoxy-D-manno-oct-2-ulosonic acid (80), or by another putative capsular polysaccharide biosynthesis proteins (81). Variants derived from clinical *Ba* isolates produce more EPS which leads to a more mucoid appearance of the colony (23), which correlates with CEP0996v and HSJ1v ability to to form biofilm than their parental strains (82) (**Figure 2A-B**). Both type of variants also produced more proteins involved in siderophore biosynthesis than their parental strain which was correlated with more siderophore production **(Figure 2A-B and C)**. pC3-null CEP0996 produces more proteases than the parental strain which could be explained by the increased abundance of other putative proteases in CEP0996 pC3-null (**Table S7;9;11**). HSJ1v lacks protease activity, likely owing to a decrease in abundance of ZmpA and ZmpB (**Figure 2D; Table S9**). Interestingly, even if higher quantity of proteins involved in the production of flagella were measured in both type of variants, CEP0996v swims more than its parental strain while this is not the case for HSJ1v (**Figure 3E**).

**Figure 2:**
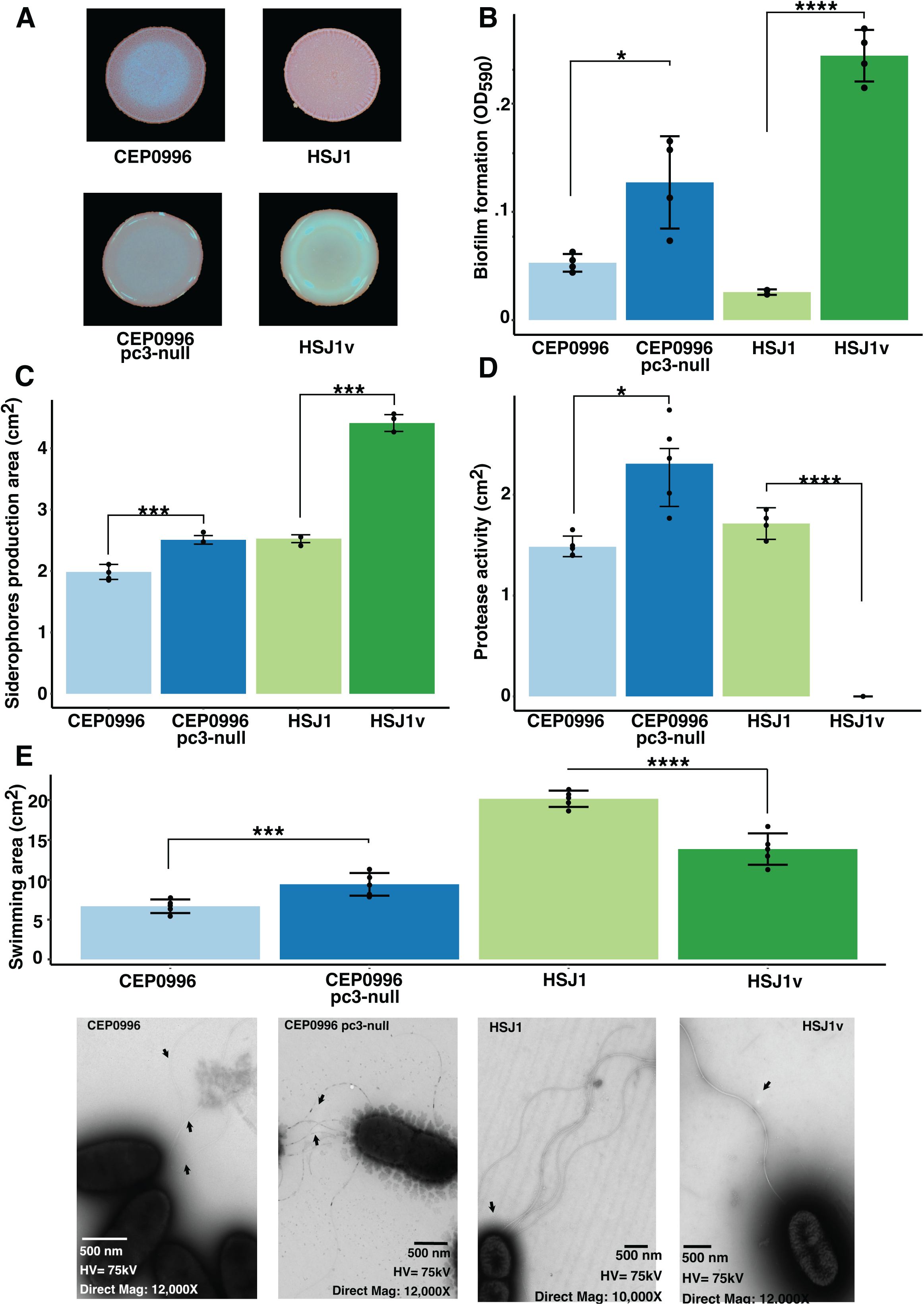
Phenotypic assays in both types of variant compared to their respective wild type. A) Morphotypes on CRTSA. B) Biofilm production (Wilcoxon test). C) Siderophore production (Wilcoxon rank test). D) Protease production (t-test). E) Swimming motility (t-test) coupled with TEM microscope of bacterium flagella. P-values are represented by * between 0.5 and 0.01, ** between 0.01 and 0.001, *** 0.001 and 0.0001, and **** inferior to 0.0001. Mean ± SD are shown on each figure.

**Figure 3:**
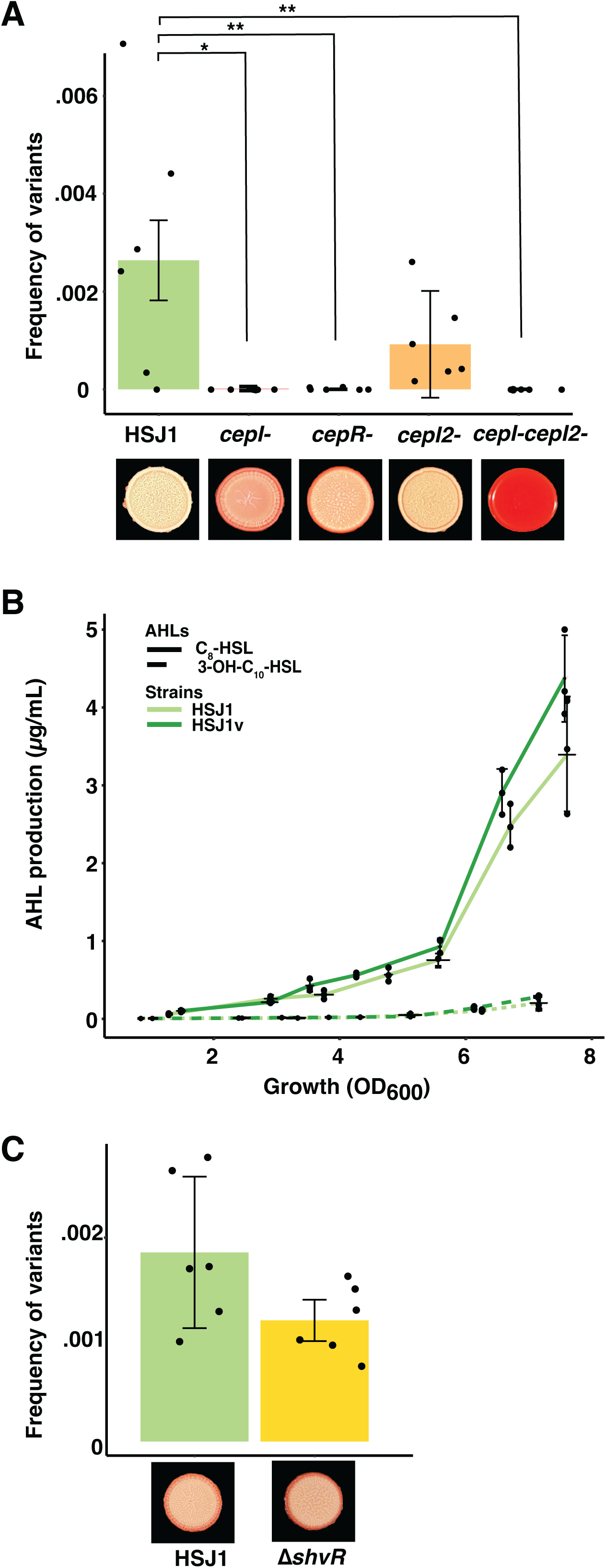
QS activates the production of phase variants while ShvR has no impact on colony morphotype and in phase variant production in *Ba* HSJ1. A) Frequency of occurrence of variant in QS mutants (Wilcoxon rank test). B) Production of AHLs in HSJ1 WT and Variant (ANOVA). C) Frequency of occurrence of variants in *ΔshvR* (t-test). P-values are represented by * between 0.5 and 0.01, ** between 0.01 and 0.001, *** 0.001 and 0.0001, and **** inferior to 0.0001. Mean ± SD are shown on each figure.

### The *B. ambifaria* HSJ1v phenotypes are not due to genomic variations

In the variant of CEP0996, the observed impact on the expression of several phenotypes is explained by loss of the pC3 replicon (32). As HSJ1v retains its pC3, we expected that the numerous phenotypic differences between WT and variant would be linked to discrete genomic differences (22). However, unexpectedly, when mapping Illumina reads of HSJ1v to the HSJ1 assembly, no significant difference was observed when results were filtered by sequencing strand bias (SPR and SAP), placement bias (EPP), variant quality (QUAL), and depth of coverage (DP) above 20 (p-value below 0.01; **Table S13**), even in the regions within the inverted-repeat sequences (**Table S14**). Furthermore, as a duplicated region was described as responsible of colony morphotype variants in *B. thailandensis* (83), we investigated if the mechanism could explain phase variation in *Ba* HSJ1. By using LASTZ, no duplicated region was found between Ba HSJ1 and its variant. These results suggests that the phase variation in HSJ1 is due to differences in modulation of gene expression or epigenetic factors rather than genomic changes.

### The *Cep* QS system impacts phase variation in HSJ1

As no genomic variation seems to be responsible for all phenotypic and proteomic differences between HSJ1 and its variant, we hypothesized that QS might, at least partially, explain some of the differences since QS regulates a large part of genes responsible for the phenotypes, including virulence and survival factors, that are different between WT and variant. In HSJ1, the Cep and Cep2 QS systems have been investigated (53, 54). *Ba* also possesses the BDSF QS system which is implicated in the regulation of c-di-GMP and thus in biofilm formation in *Bc* (55–59). In our proteomic study of HSJ1 at OD_600_ ∼1, proteins CepI and CepR were detected, as well as RpfF and RpfR belonging to the BDSF QS system. Neither CepI2 nor CepR2 were detected (**Table 2; Table S9**).

**Table 2.** Quorum sensing-related gene/proteins detected in the proteomic study of strains HSJ1.

|  | Locustag | Detection<br>in<br>proteomic<br>analysis | Quantifiable in WT | Quantifiable in variant |
| --- | --- | --- | --- | --- |
| *cepl* | BAMB_HSJ1_RS20835 | + | $2.82 \cdot 10^8 \pm 1.05 \cdot 10^7$ | $1.47 \cdot 10^8 \pm 9.02 \cdot 10^6$ |
| *rsaM* | BAMB_HSJ1_RS20840 | + | $1.35 \cdot 10^8 \pm 2.80 \cdot 10^7$ | $1.2 \cdot 10^8 \pm 3.29 \cdot 10^7$ |
| *cepR* | BAMB_HSJ1_RS20845 | + | $2.27 \cdot 10^7 \pm 1.15 \cdot 10^7$ | $3.62 \cdot 10^7 \pm 7.57 \cdot 10^6$ |
| *cepl2* | BAMB_HSJ1_RS28550 | + | $5.62 \cdot 10^7 \pm 1.47 \cdot 10^7$ | $2.01 \cdot 10^7 \pm 1.59 \cdot 10^6$ |
| *cepR2* | BAMB_HSJ1_RS28485 | - | NA |  |
| *rpfF* | BAMB_HSJ1_RS26685 | + | $5.10 \cdot 10^7 \pm 1.68 \cdot 10^7$ | $3.46 \cdot 10^7 \pm 1.10 \cdot 10^7$ |
| *rpfR* | BAMB_HSJ1_RS26690 | + | $2.81 \cdot 10^8 \pm 1.14 \cdot 10^8$ | $3.61 \cdot 10^8 \pm 4.65 \cdot 10^7$ |
| *shvR* | BAMB_HSJ1_RS29045 | + | $5.19 \cdot 10^8 \pm 2.00 \cdot 10^8$ | $0 \pm 0$ |

Values based on LFQ Intensities, after filtering and imputing missing values

To evaluate the impact of QS *per se* on phase variation, we determined the frequency of occurrence of variants in HSJ1 and its isogenic *cepI-, cepR-, cepI2-,* and *cepI-cepI2-* mutants. Interestingly, the *cepI-, cepR-,* and *cepI-cepI2-* did not produce any variants while *cepI2-* produced variants statistically at the same frequency as the parental strain (**Figure 3A; Table S15**). When complemented with a plasmid expressing the *cepR* gene, the *cepR*-mutant recovered its ability to generate variants (**Figure S3**), confirming that the CepI/CepR QS system is involved in the occurrence of phenotypic variants.

As we previously reported (23), no difference in production of both AHLs between HSJ1 and its variant was observed (**Figure 3B**). Due to the loss of pC3 where *cepI2* is located, CEP0996v does not produce 3-OH-C_10_-HSL while there is no difference in C_8_-HSL production between CEP0996 and its variant (**Figure S4**). Thus, while QS seems important for the emergence of variants, phase variation does not apparently impact the function of AHL-dependant QS.

The protein ShvR regulates QS and impacts colony morphology, virulence, and biofilm formation in the *Bc* strain K56-2 (27, 42). We hypothesize that the protein encoded by gene BAMB_HSJ1_RS29045 (87.19% identity with the *shvR* gene of *Bc* and with 90.03% similarity at protein level; **Figure S5**) could be the ShvR homologue in *Ba* HSJ1 with a function similar to the one described in *Bc* K56-2. Interestingly, this ShvR homologue is lost in HSJ1v compared to HSJ1 (**Table 2; Table S9**). However, deleting BAMB_HSJ1_RS29045 in HSJ1 did not impact the occurrence of variants (**Figure 3C; Table S16**). Furthermore, colony morphology and biofilm formation were not affected by the deletion BAMB_HSJ1_RS29045 in HSJ1 (**Figure S6**). These data strongly suggest that this protein has a different function in HSJ1 than in *Bc* K56-2. Intriguingly, inactivation of the *shvR* homologue in the *Bc* strain H111 was also unaltered (44).

Altogether, these results show that the CepI/CepR QS system is important for the emergence of phase variants in HSJ1, but not ShvR.

### DNA methylation inhibits the emergence of phase variants

QS is thus involved in the occurrence of variants in a *Ba* strain which does not rely on loss of pC3. Since no genetic modifications or impaired QS could apparently explain the major differences in phenotypes observed between HSJ1 and its variant, we hypothesized that an epigenetic factor such as DNA methylation could be playing a role in phase variation, as it was found to affect a few similar phenotypes in *Bc* (47). Indeed, DNA methylation is a recognized mechanism driving phase variation is several bacteria (e.g. *Neisseria*, *Helicobacter pylori*) (84)).

Three adenosine DNA methylation motifs were identified in the HSJ1 genome using PacBio sequencing which are homologs to the identified adenosine DNA MTases in *Ba* strain AMMD (46) (**Table 3**). No SNP in the three DNA MTase sequences were found between variant and WT (**Figure S7**).

**Table 3.** DNA methylation in strain HSJ1.

| Motif | Methylation | Strand | Strain | Called modified motifs* | % Motif Detected | Mean Modif. QV | Mean Motif Coverage | Putative Genes and their putative homologs in other Bcc |
| --- | --- | --- | --- | --- | --- | --- | --- | --- |
| CACAG | m6A | F | HSJ1 | 6265/6327 | 96.12 | 70.03 | 39.72 | BAMB_HSJ1_RS01220 - type III (DNA Mtase 1) |
|  |  |  | HSJ1v | 6260/6327 | 98.94 | 72.94 | 41.87 |  |
| GTWWAC | m6A | F+R | HSJ1 | 1578/1680 | 93.92 | 62.86 | 39.61 | BAMB_HSJ1_RS24645 - type II orphan (DNA Mtase 2) |
|  |  |  | HSJ1v | 1599/1680 | 95.17 | 65.13 | 41.7 |  |
| RGATCY | m6A | F+R | HSJ1 | 11402/11862 | 96.12 | 68.34 | 39.76 | BAMB_HSJ1_RS24550 - type II (DNA Mtase 3) |
|  |  |  | HSJ1v | 11501/11862 | 96.95 | 69.83 | 41.87 |  |

In our proteomic analysis, two out of the three corresponding DNA MTase (1 and 2) proteins were detected in both the parental strain and the variant at similar levels at OD_600_ of ∼1, ∼3, and ∼5 while DNA MTase 3 was not detected (**Figure S8**; **Table S9**). The three DNA MTase-encoding genes were deleted to investigate their impact on phenotypic variation in HSJ1 (**Figure 4; Table S17-18**). Supporting the hypothesis of an epigenetic factor regulating the emergence of variants, the frequency of occurrence of phase variation was significantly increased in the DNA MTase 2 mutant (responsible for the GTWW**^6^<u>A</u>**C motif) and was lowered when the gene encoding for this DNA MTase was overexpressed in the mutant background (**Figure 4; Table S17**).

**Figure 4:**
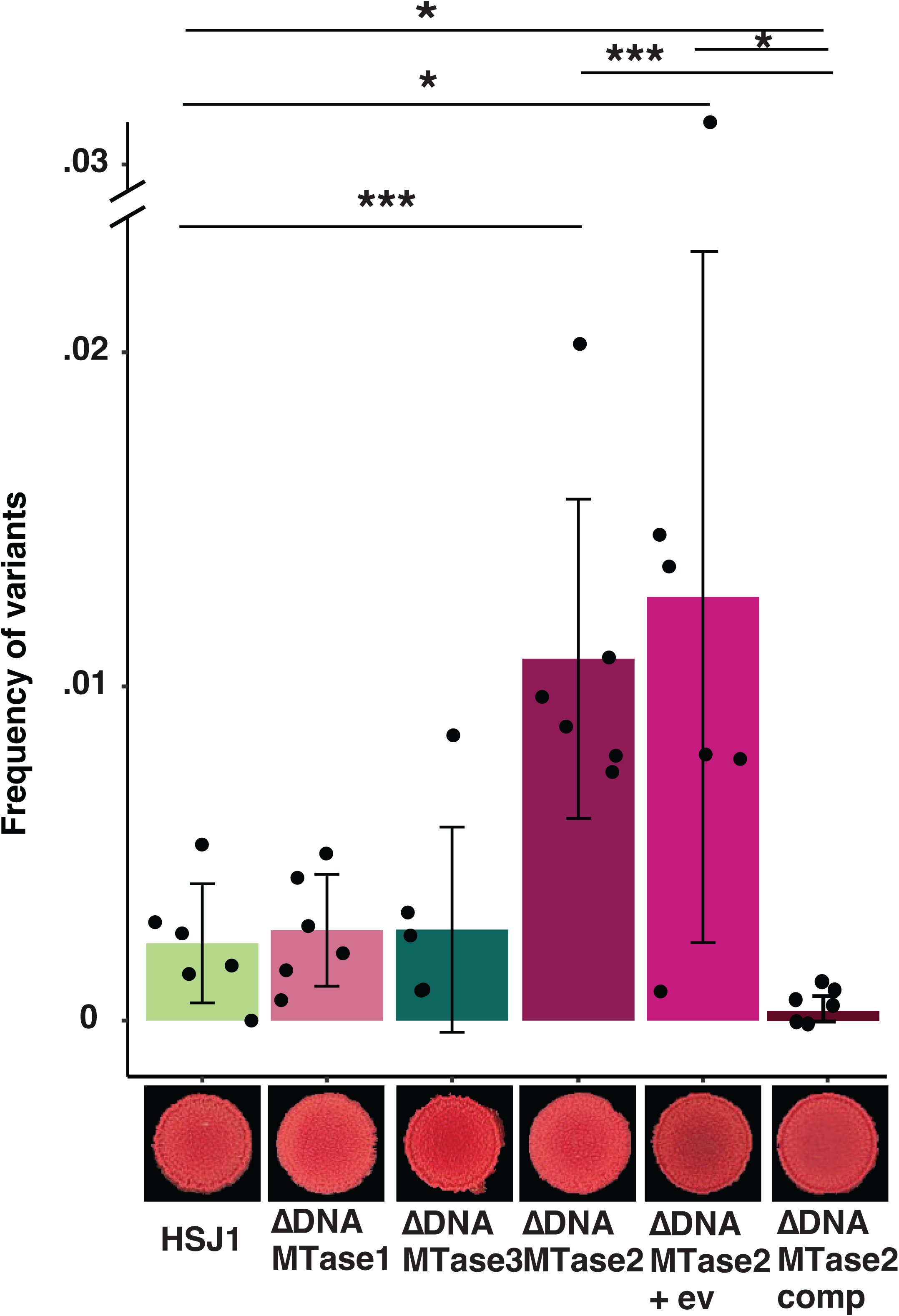
Impact of adenosine DNA MTases in *B. ambifaria* HSJ1 on phase variation. Frequency of occurrence of variant in the three DNA MTases mutants and colony morphotype on CRLA (Wilcoxon rank test). P-values are represented by * between 0.5 and 0.01, ** between 0.01 and 0.001, *** 0.001 and 0.0001, and **** inferior to 0.0001. Mean ± SD are shown on each figure. “+ ev” means empty vector, “comp” means deletion was complemented with a constitutive promotor.

DNA methylation – through the DNA MTase 2 – appears to be involved in the emergence of the phase variants by inhibiting a still unknown mechanism. Based on phenotypic assays (**Figure S9**), there is no difference in biofilm formation between HSJ1 and Δ*DNAMTase 2*, however the constitutive transcription of DNA MTase 2 significantly increases biofilm formation (**Figure S9A**). Deleing the DNA MTase 2 gene does not lead to difference in protease activity (**Figure S9B**). Also, ΔDNA MTase 2 mutant is less motile than the WT probably because of higher proportion of variants (**Figure S9C**). DNA MTase 2 has no impact on swimming motility and protease production, but rather the production of variant in the mutant has an impact on observed phenotypes (**Figure S9B-C**).

### QS and DNA methylation do not interplay

With QS and DNA MTase 2 having opposite effects on the emergence of HSJ1 variants, one could influence the other. To answer this question, we first quantified the abundance of the DNA MTase 2 protein at OD_600_∼3 and ∼5 in the HSJ1 *cepR-* QS mutant compared to HSJ1 WT. No different in abundance was measure (**Figure S10A; Table S19**). This is compatible with the apparent absence of a cep box (54) in the promoter region of the BAMB_HSJ1_RS_24645 gene, encoding for DNA MTase 2. Then, we quantified the abundance of proteins belonging to Cep QS, Cep2 QS and Hmq systems as well as their respective signal molecules: C_8_-HSL, 3-OH-C_10_-HSL and HMAQ-C_7_:2’ (**Figure 5; Tables S19-20**) as these systems are known to interplay (53). While no significant difference in abundance of CepI, RsaM, CepR, CepR2 and HmqA were observed between HSJ1 Δ*DNAMTase2* and HSJ1 WT, CepI2 was at lower abundance than the detection level by LC-MS/MS at OD_600_∼3 but was quantified at similar abundance at OD_600_∼5. However, the level of 3-OH-C_10_-HSL and HMAQ-C_7_:2’ were reduced in HSJ1 ΔDNAMTase2 at OD_600_∼3 and ∼5 (**Figure 5; Figure S10**). This result is not unexpected as we have shown that the DNA MTase2 mutant produces more variants than the WT (**Figure 4**) and variants of HSJ1 do not produce HMAQs, such as HMQ-C_7_:2’, at all (23).

**Figure 5:**
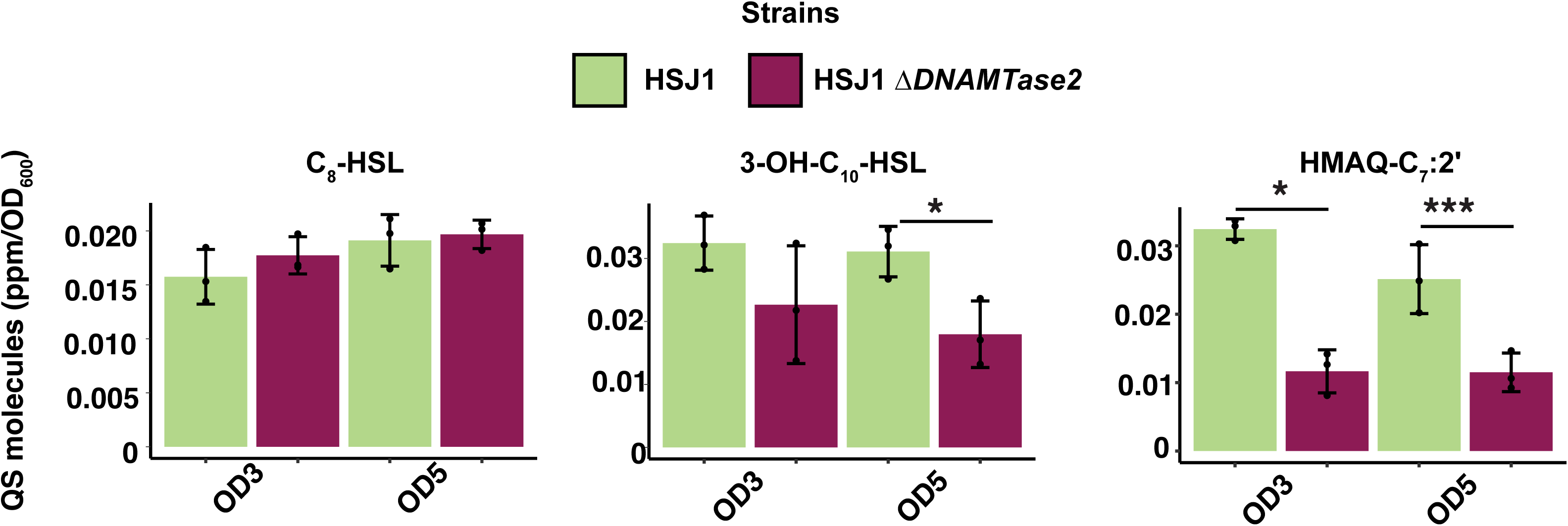
DNA MTase 2 impairs signal molecules produced by Cep2 and Hmq systems, at OD_600_ 3 and 5. Production of AHLs and HMAQ in HSJ1 ΔDNA MTase 2 mutant (t-test). P-values are represented by * between 0.5 and 0.01, ** between 0.01 and 0.001, *** 0.001 and 0.0001, and **** inferior to 0.0001. Mean ± SD are shown on each figure.

## Discussion

Bcc infections are acquired from the environment with some patient-to-patient transmission (85–88). During long-term infection in CF, *Burkholderia* uses phase variation to persist and adapt to changes within the human host by decreasing the production of virulence and immunogenic factors (73, 89). In members of the Bcc, virulence is affected by their ability to [1] produce quorum-sensing signal molecules, [2] modulate virulence factors such as siderophores, proteases and biofilm formation, [3] resist to antibiotics and [4] infect and replicate within macrophages. An increased resistance to antimicrobial agents, and decreased biofilm formation, loss of mucoidy, and reduced motility are well-established to play a role in *Burkholderia* persistence during infection (34, 35, 73, 90). This variation in morphology and phenotypes, within patients, is a natural mechanism and reversion or modulation are observable under laboratory conditions (23–25, 28, 29, 32, 33, 63, 64, 91–93).

Previously described clinical *Ba* strains undergoing phase variation, while sharing similarities in modulation of virulence and survival factors (23), do not use the same mechanism. As observed in other species belonging to the Bcc (32, 43–45), some *Ba* variants have lost their pC3 which explains most of the observed changes in phenotypes in those variants (46). Based on phenotypic and proteomic assays, *Ba* HSJ1 WT is closest to small colony variant and Ba HSJ1v to large colony variant described for *Bp* (62–65). However, other clinical *Ba* variants keep their pC3 and the emergence of variants is impacted by both QS and DNA methylation (**Figure 6**). The mutation of both systems does not explain all phenotypic differences observed between parental strain and variant. Thus, it is more likely that both systems regulate either the promotor region of the genes encoding for virulence factors or a general transcription regulator. In *Bc* K56-2, ShvR is known to regulate QS and virulence factors, but we showed that the homolog and ortholog of this LysR-type transcriptional regulator does not have the same function in *Ba* HSJ1 – a similar observation that was made in Bc H111 (44). In *Bp*, YelR (Yellow program regulator) was identified to play a role in colony morphotype variation by producing smaller smooth colony compared to bigger rough colonies. In HSJ1, BAMB_HSJ1_RS_07310 is an ortholog and homolog (86% identity) to YelR and could be a potential candidate (25). In *B. thailandensis*, a secondary metabolite regulator (ScmR) LysR-type transcriptional regulator has been shown to control various biosynthesis gene clusters, biofilm formation and virulence as well as QS (94, 95). In HSJ1, BAMB_HSJ1_RS_03645 is an ortholog and homolog (76% identity) to ScmR and could be a potential candidate for further studies to determine the mechanism underlying phase variation in HSJ1.

**Figure 6:**
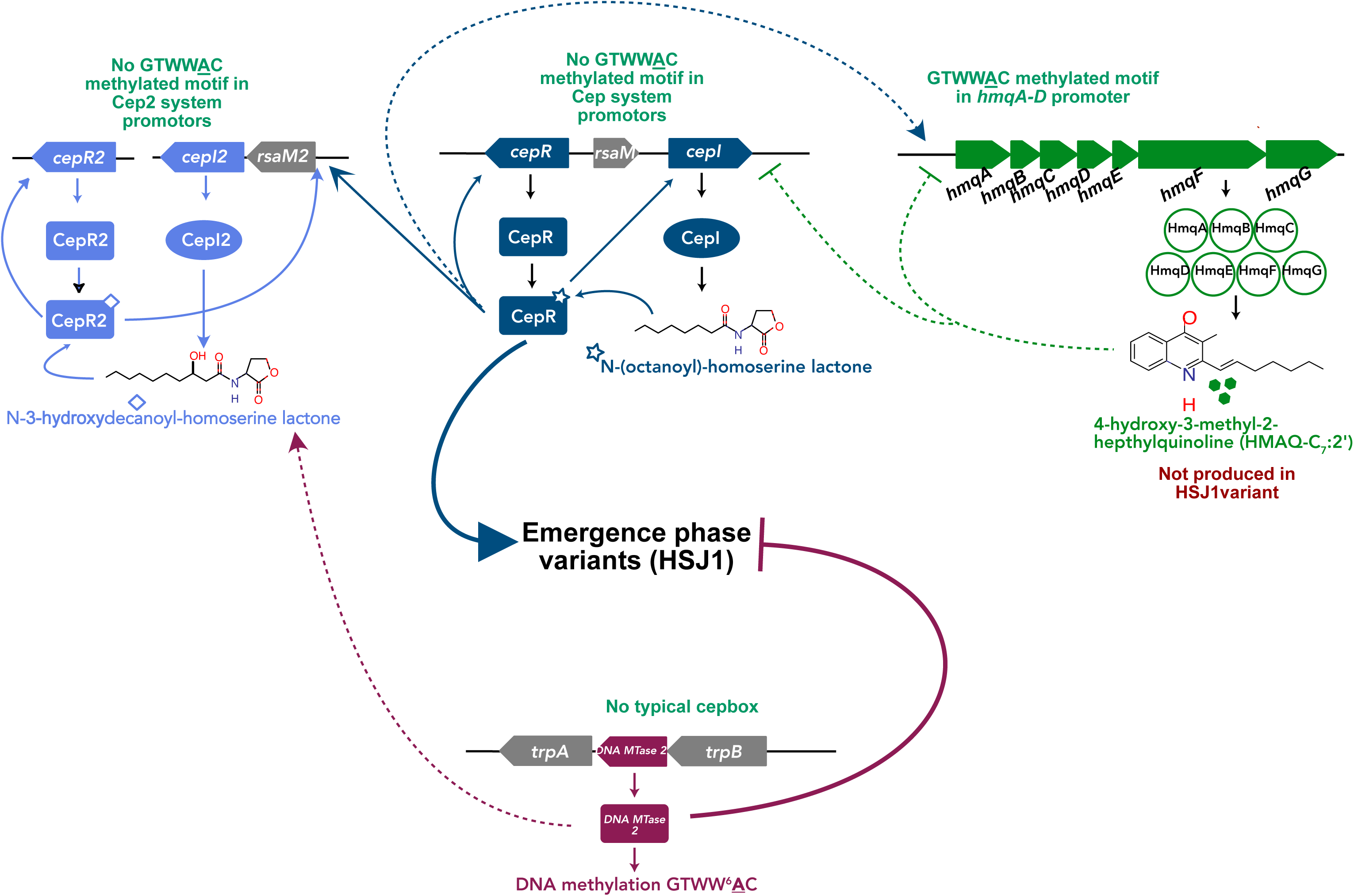
Schematic representation of phase variation in *Ba* HSJ1 and the roles of both AHL-based QS systems, Hmq system and DNA MTase 2. Cep QS system activates phase variation in *Ba* HSJ1 while DNA MTase 2 inhibits the emergence of variants. Cep QS system does not regulate the production of DNA MTase 2 proteins. While the DNA MTase 2 does not significantly impact the abundance of proteins involved Cep, Cep2 and Hmq systems, the production of both 3-OH-C_10_-HSL and HMAQs were reduced due to the production of variant.

Only a few studies reported *Burkholderia* colony morphotype variation and most of them did not report the underlying mechanism. As colony morphotype variation is prevalent during infection and impact virulence, it could play a crucial role in *Burkholderia* persistence during infection. Before investigating the multiple mechanisms underlying phase variation in *Burkholderia*, a comprehensive study determining the prevalence of the colony morphotype variation phenomenon among diverse ecological niches of *Burkholderia* is necessary. Understanding *Burkholderia* colony morphotype variation and the underlying mechanisms will enhance the development of more efficient antibiotic treatments and vaccines.

## Material and Methods

### Screening for the presence of the third replicon in wild type and variants

All strains, and plasmids are listed in **Tables S1** and **S2**, respectively. Overnight cultures were set up from -80°C glycerol stocks in Tryptic Soy broth (TSB, Bacto BD Difco) at 30°C with shaking at 200 rpm. DNA was extracted from cultures following a previously described method using a Fast-Prep Beadbeater (96). PCR amplification was performed using the Easy-Taq DNA polymerase (Transgen) with specific primers targeting the *hisA* gene, which is present on the first chromosome, or seven different regions distributed along pC3 in *Ba*. All primers are listed in **Table S3**.

### Whole Genome Sequencing and representation

*Ba* strains CEP0996 and HSJ1 and their respective phase variants were cultured overnight from -80°C-stored glycerol stocks in TSB at 30°C with shaking at 200 rpm. Genomic DNA (gDNA) was extracted from cultures using a Genomic DNA extraction kit (Transgen). *Ba* strains CEP0996 and HSJ1 and their respective variant’s genomes were sequenced using either Oxford Nanopore Technology (ONT, by MIGS) or Pacific Biosciences technology (PacBio, by Genome Quebec) RS II complemented by Illumina technology (MIGS).

HSJ1 WT and variant genomes were assembled using Unicycler (conda v0.4) hybrid method (97), from both long and short reads. For CEP0996 WT, Unicycler hybrid (docker v0.5), Canu v0.0 (98) and Flye v2.9.3-b1797 (99) assemblies were used to generate a consensus genome assembly using Trycycler v0.5.4 (100). Then, Medaka v1.0.3 from ONT and Polypolish v0.5.0 (101) tools were used to polish the resulting Trycycler assembly. Each genome assembly was then annotated using BAKTA v1.9.1 (102). To confirm the absence of the pC3 replicon in both type of variants, short reads from *Ba* CEP0996 pC3-null and *Ba* HSJ1v were mapped using minimap2 (103, 104) on their corresponding WT genomes. Coverage of reads for each chromosome was represented using one in every 100 nucleotides using Shinycircos R package (shiny::runGitHub ("shinyCircos", "venyao"); (105)).

To assign homolog genes between CEP0996 and HSJ1, a megablastn was set up with only one output match for each gene in query using the following parameters: - evalue 10 -outfmt 6 -num_threads 3 -max_target_seqs 1 -max_hsps 1 (106).

### Comparison between genomes of HSJ1 and HSJ1 variant

Illumina reads from *B. ambifaria* HSJ1 variant (HSJ1v) were mapped on the *B. ambifaria* HSJ1 WT genome assembly, and *B. ambifaria* HSJ1 WT Illumina reads were mapped on *B. ambifaria* HSJ1 variant genome assembly using minimap2.1 (103, 104). Then, sorted mapped reads were used by Freebayes v1.3.2-dirty (107) to detect any genomic variations between HSJ1 WT and HSJ1v using diploid feature.

Both HSJ1 and HJS1v assembly were screened for inverted repeat sequences using Inverted Repeats finder tool with default settings (https://tandem.bu.edu/irf/home; (108)) and for duplication regions using LASTZ software (https://github.com/lastz/lastz; (109)).

### Comparative proteomic between both CEP099 WT/variant and HSJ1 WT/variant couples

Colonies were isolated on tryptic soy agar plates containing 0.01% Congo Red (CRTSA) following an incubation at 30°C for 48 h. Cultures for both WT and variants of CEP0996 and HSJ1 were prepared in TSB and incubated overnight at 30°C with agitation in a TC Roller-drum (New Brunswick) at 200 rpm. Fresh cultures were set up at an OD_600_ of 0.2 in 5 mL of TSB until the OD_600_ reached 1. Total protein was extracted as previously described (110). Briefly, after washing cell pellets with phosphate-buffered saline (PBS), frozen cells were lysed, and total protein was precipitated with 80% ice-cold acetone.

Twenty µg of each sample were reduced and alkylated by 0.2 mM DTT and 0.8 mM iodoacetamide. Then samples were digested with 0.4 µg trypsin and purified on C18 StageTips, and 1 µg analysed following a 120 min liquid chromatography run coupled with mass spectrometry (LC-MS/MS) on an Orbitrap Fusion Tribrid system (Thermo Fisher) using DDA label-free technique at the Proteomic Platform of the CHU Québec Research Center (Université de Laval, Quebec City, Canada). Raw data were searched either against CEP0996 or HSJ1 proteomes, obtained from this study annotated genomes, (**Table S4-5**) by Fragpipe v20 (111) using the LFQ-MBR script for label-free quantification of proteins and normalization. The resulting “combined proteins” files were filtered to remove contaminants and keep proteins for which there were at least two values for at least one group based on the LFQ intensities as each data search contains three biological replicates for each condition. Data was normalized using the variance stabilization normalization (112) then selected to differentiate proteins with at least two missing values in one condition representing a “presence/absence”. Data containing one missing value in at least one group had the missing value imputed based on the average of the values of other replicates within the same group and the differential protein production between each group was analysed by using a Welch’s t-test with Benjamini-Hochberg adjusted pvalues below 0.05 (113, 114). Analyses were performed on three biological replicates.

### Phenotypic Assays

Colony morphologies were observed by spotting 15 µL from a TSB overnight culture on CRTSA plates. Plates were incubated for two days at 30°C.

Biofilm formation was measured by growing static cultures in 5 x 75-mm polystyrene tubes or in 96-well plates. Cultures were inoculated from an overnight culture to an initial OD_600_ = 0.2 and were incubated at 30°C for 24 h without agitation. After removing unattached cells by washing, the biofilms were stained with 0.1% crystal violet and OD_590_ was measured using a microplate plate reader (Cytation 3, BioTek or Clariostar) (23). Four technical replicates were set up for the 96-well plate experiments and four biological replicates were used for biofilm formation in tubes. The experiment was performed three times.

Siderophore production was determined by Chrome-Azurol S (CAS) assay (115). A 5 µL bacterial suspension at OD_600_ = 5 was spotted on CAS agar plates. Plates were incubated overnight at 30°C. Areas of the siderophore production halos surrounding the colonies were then measured and reported in cm^2^. Five biological replicates were used, and the experiment was performed three times.

Protease activity was determined by spotting 15 µL of a bacterial culture at OD_600_ = 5 on 1.5% skim milk TSA plates, as previously described (54). Plates were incubated 48 h at 30°C. Protease activity halos were then measured and reported in cm^2^. Four biological and three technical replicates were used, and the experiment was performed three times.

Bacterial flagella were observed by transmission electron microscopy (TEM). CEP0996 strains were cultured for 48 h on CRTSA plates and HSJ1 strains were cultured in 5 mL TSB at 30°C overnight with agitation. A grid was inoculated into the liquid culture for 30 seconds and dried for 1 min. Then, cells were fixed for one second using a 1% paraformaldehyde solution (PFA) before imaging using a Hitachi H-7100 electron microscope (INRS-CAFSB) platform with AMT Image Capture Engine (version 600.147).

### Determination of phase variation frequency

From single colonies, starter cultures were set up in TSB and incubated at 30°C overnight under 200 rpm shaking. The next morning, 15 µl of each culture was spotted on a LB agar plate containing 0.01% Congo Red (CRLA) and incubated for 48 h at 30°C followed by 96 h at room temperature. Each spot colony was then suspended in 1 mL PBS and serially diluted. Finally, 50 µL of the 10^6^ dilution was spread onto a CRLA plate. Three sets of dilutions were performed and plated from each colony. After incubation at 30°C, variants were counted among the total colonies per plate to determine the frequency of occurrence of variant colonies per cell per generation (116). The experiment was performed on six separate colonies and repeated twice.

### Comparative proteomics HSJ1, HSJ1 Δ*DNA MTase 2*, HSJ1 *cepR-* strains

Concerning samples preparation for proteomics to compare abundance of protein of interest between HSJ1 WT, HSJ1 *cepR*- and HSJ1 ΔDNAMTase2, samples were cultured as previously described until the OD_600_ reached ∼3 and 5. Total proteins were extracted by lysing the cells in 1% SDC in 100 mM HEPES pH 8.5. Then, proteins were quantified using a BCA assay and 20 µg proteins were trypsin-digested and cleaned on one SBD-RPS disk to normalized proteins to 10 µg for each sample. Using an Acquity M-class nanoLC system (Waters, USA), 5 µL of the sample was loaded at 15 µL/min for 3 minutes onto a nanoEase Symmetry C18 trapping column (180 µm x 20 mm) before being washed onto a PicoFrit column (75 µm x 350 mm; New Objective, Woburn, MA) packed with SP-120-1.7-ODS-BIO resin (1.7 µm, Osaka Soda Co, Japan) heated to 45°C. Peptides were eluted from the column and into the source of a Q Exactive Plus mass spectrometer (Thermo Scientific) using the following program: 5-30% MS buffer B (98% Acetonitrile + 0.2% Formic Acid) over 90 minutes, 30-80% MS buffer B over 3 minutes, 80% MS buffer B for 2 minutes, 80-5% for 3 min. The eluting peptides were ionised at 2400V. A Data Dependant MS/MS (dd-MS2) experiment was performed, with a survey scan of 350-1500 Da performed at 70,000 resolution for peptides of charge state 2+ or higher with an AGC target of 3e6 and maximum Injection Time of 50ms. The Top 12 peptides were selected fragmented in the HCD cell using an isolation window of 1.4 m/z, an AGC target of 1e5 and maximum injection time of 100ms. Fragments were scanned in the Orbitrap analyser at 17,500 resolution and the product ion fragment masses measured over a mass range of 120-2000 Da. The mass of the precursor peptide was then excluded for 30 seconds. Raw Data search was done against HSJ1 proteome without match between runs. Data analysis was proceeded as described above using LFQ intensities.

### Quantification of signal molecules

AHLs and HMAQs were extracted from 4 mL cultures with ethyl acetate (1:1) and concentrated 10 times in acetonitrile, as previously described (53, 54, 117). 5,6,7,8-tetradeutero-4-hydroxy-2-heptylquinoline (HHQ-d4) was used as an internal standard. Samples were analysed by liquid chromatography coupled with mass spectrometry (LC-MS) in positive electrospray ionization using a Kinetex 5-m m EVO C18 100-Å 100-by 3-mm reverse-phase column. A Quattro Premier XE triple quadrupole was used as the detector (Waters). A multiple reaction monitoring (MRM) program was used to detect HMAQ families. This experiment was conducted with three independent biological replicates.

### DNA methylome of *B. ambifaria* HSJ1 WT

Blasr (118), samtools (119), kinetictools and motifmaker-master, bax2bam, pbcoretools from PacBio github (https://github.com/PacificBiosciences) tools were used to determine the methylome of HSJ1 WT from PacBio data sequencing based on Ba HSJ1 genome assembly (120, 121).

### Mutant construction in *B. ambifaria* HSJ1

Single mutants of BAMB_HSJ1_RS01220 (CAC**<u>A</u>**G motif named as DNA MTase 1), BAMB_HSJ1_RS24645 (GTWW**<u>A</u>**C motif named as DNA MTase 2) and BAMB_HSJ1_RS24550 (RG**<u>A</u>**TCY motif named as DNA MTase 3) putative DNA MTase-encoding genes and BAMB_HSJ1_RS29045 (putative *shvR*), were generated using homologous recombination with the pEX18Tet-PheS allelic replacement plasmid (122). Sequences of 5’UTR and 3’UTR both including nine nucleotides of the encoding sequence of targeted genes and trimethoprim resistance gene were amplified with flanking regions by PCR using specific primers (**Table S3**) and polymerase (EasyTaq [Transgen, China] or Phusion HiFi polymerase [Thermofisher, Australia]). PCR-amplified inserts and the pEX18Tet-PheS previously digested by BamHI and PstI (NEB, Australia or Thermofisher, Canada) were assembled using the Seamless Cloning method and transformed into *E. coli* Trans-T1 or *E. coli* NEB5ɑ. The resulting plasmids were transformed into conjugative *E. coli* SM10 to transfer each plasmid into HSJ1 using bi-parental conjugation. Then, tetracycline-resistant recombinants were selected on TSA with 250 µg/mL tetracycline and spread on M9 agar plates containing 100 µg/mL trimethoprim and 0.1% chlorophenylalanine (Sigma, USA) to cure pEX18Tet-PheS. Resulting secondary recombinants were transformed with pFLe4 (30°C) plasmid on LB agar plate with 300 µg/mL kanamycin - to flip the trimethoprim resistance gene. Resulting colonies were double screened on LB agar plate containing 250 µg/mL trimethoprim to select trimethoprim sensitive mutants (**Table S1**).

### Complementation of DNA MTases genes in *Ba* HSJ1

Each putative DNA MTase-encoding gene was amplified using EasyTaq (Transgen, China) or Phusion HiFi polymerase (Thermofisher, Australia) and KpnI and HindIII restriction sites were added in the primers used. The pMLS7 plasmid encoding for a constitutive S7 promotor was digested by KpnI and HindIII enzymes (NEB, Australia). Both resulting PCR fragment and digested plasmid were purified on an agarose gel and ligated using the T4 DNA ligase (NEB, Australia). The resulting DNA MTases complementation vectors were transformed into *E. coli* SM10 and were transferred by bi-parental conjugation into the different *Ba* HSJ1 DNA MTase mutants (**Table S1**).

## Data availability

Genomes from this study are available in GenBank under the Bioproject: PRJNA896258 (123, 124). Proteomic raw data deposited to the ProteomeXchange Consortium via the PRIDE partner repository with the dataset identifier PXD055955 (125).

## Acknowledgements

The authors would like to thank Dr Piklu Bhattacharya for her insight in genomic assemblies; Julien L. Breton-Robin for helping setting command lines tools, especially writing a script used for the methylome analyses and Dr Max Cummins for his assistance in usage of high-performance computing resources used for this research, as well as for reviewing the article.

This work was supported by grants MOP-142466 from the Canadian Institutes of Health Research (CIHR), PRO24-19185 from the University of Technology 2024 Early Career Researcher Capability Development Initiative, and Science Seed funding from the University of Technology Sydney and from the National Health and Medical Research Council of Australia (GNT1194325). E.D. held the Canada Research Chair in Sociomicrobiology.

P.M.L.C.: conception, design of the manuscript and acquisition of data, interpretation of data, and revision of the manuscript; M-C.G: acquisition of data, interpretation of data, and revision of the manuscript; A.H.: revision of the manuscript, T.P.S.: resources, funding and revision of the manuscript; M.P.P: proteomics data analysing troubleshooting, revision of the manuscript; E.D.: resources, funding, interpretation of data, and revision of the manuscript.

